# Wide-field bond-selective fluorescence imaging: from single-molecule to cellular imaging beyond video-rate

**DOI:** 10.1101/2024.07.05.601986

**Authors:** Dongkwan Lee, Haomin Wang, Philip A. Kocheril, Xiaotian Bi, Noor Naji, Lu Wei

**Affiliations:** Division of Chemistry and Chemical Engineering, California Institute of Technology, Pasadena, California 91125, USA

## Abstract

Wide-field (WF) imaging is pivotal for observing dynamic biological events. While WF chemical microscopy offers high molecular specificity, it lacks the sensitivity for single-molecule detection. In contrast, WF fluorescence microscopy provides live-cell dynamic mapping but fails to leverage the rich chemical information necessary for functional interpretations. To address these limitations, we introduce Wide-Field Bond-selective Fluorescence-detected Infrared-Excited (WF-BonFIRE) spectro-microscopy. This technique combines rationally optimized imaging speed and field-of-view (FOV) to achieve single-molecule sensitivity with bond-selective contrast. WF-BonFIRE outperforms its point-scanning counterpart, enhancing frame acquisition up to 10,000 times. We demonstrate WF-BonFIRE’s capabilities in imaging cells, astrocytes, and live neurons, capturing single FOVs up to 50 µm × 50 µm, with further expansion via multi-FOV mosaicking. Additionally, we have implemented a temporal-delay modulation scheme that allows real-time kilohertz imaging speeds up to 1500 Hz. This enables millisecond temporal resolution while monitoring random motion of live Escherichia coli. Overall, WF-BonFIRE significantly broadens the possibilities for chemical imaging, enabling high-speed observations at unparalleled sensitivity levels.

**One-Sentence Summary:** Wide-field bond-selective fluorescence imaging pushes chemical-sensitive microscopy platform into a new regime, achieving single-molecule sensitivity and speeds up to kilohertz.

## INTRODUCTION

Biological processes are inherently heterogeneous and dynamic. Consequently, the ability to image subcellular live-cell activities with high-sensitivity and fast speed has revolutionized our understanding of fundamental biology (*1, 2*). To this end, point-scanning microscopy configurations face fundamental constraints to achieve high temporal resolution due to serial pixel-by-pixel acquisition (*1, 3, 4*). In contrast, wide-field (WF) microscopy, with down to single-molecule sensitivity, enables the capture of dynamic biomolecular processes, such as those involving RNA, proteins, and metabolites, at video-rate speeds across the entire field of view (FOV). This capability is particularly valuable for applications like imaging and tracing neuronal action potentials that occur at millisecond timescales across entire neurons and neighboring cells (*2, 5*). Furthermore, leveraging the high-sensitivity capabilities of wide-field imaging, advanced functional fluorescence microscopy techniques have been developed, including single-molecule localization microscopy (SMLM), single-molecule fluorescence in situ hybridization (smFISH) microscopy, and structured illumination microscopy (SIM), which have significantly advanced our understanding of complex and dynamic biological phenomena to unprecedented levels (*6*–*9*).

Over the past two decades, chemical imaging techniques have dramatically advanced, offering high molecular specificity and detailed bond-selective information crucial for biological research (*10*). Despite these advances, achieving single-molecule sensitivity in wide-field (WF) chemical imaging remains a formidable challenge. Raman scattering suffers from inherently small cross-sections of vibrational transitions, ranging from 10^−30^ to 10^−28^ cm^2^, more than ten orders of magnitude smaller than the visible absorption cross-sections in fluorescence spectroscopy (*11*). Nonlinear Raman microscopy, such as stimulated Raman scattering (SRS), addresses this limitation with up to 10^8^ stimulated emission amplification with two simultaneous pulsed lasers (*12*). However, the reliance on tightly-focused nonlinear excitation limits the scope of WF biological imaging, even with the most recent advances of electronic pre-resonance SRS (epr-SRS) and stimulated Raman-excited fluorescence (SREF) (*13*–*16*). Conversely, mid-infrared (MIR) spectroscopy, with its substantially larger linear MIR absorption cross-sections (10^−22^ – 10^−17^ cm^2^), enhances the feasibility of high-speed WF imaging (*17*–*19*). Recent breakthroughs in MIR photothermal microscopy, for instance, have achieved WF capabilities for ultrafast imaging (*20*– *22*). However, the sensitivity of these methods is still confined to the micromolar to millimolar range, leaving the pursuit of high-sensitivity WF chemical imaging—from low micromolar down to single-molecule detection—for more universal biological targets largely unmet.

Recently developed bond-selective fluorescence-detected infrared-excited (BonFIRE) microscopy has advanced to achieve single-molecule sensitivity while capturing mid-infrared chemical information for live biological imaging, representing a significant leap in bio-imaging capabilities (*23, 24*). Employing a picosecond MIR and near-infrared (NIR) double-resonance excitation scheme, BonFIRE transfers the rich MIR-excited vibrational information into fluorescence signals for highly sensitive detection **(Fig. 1A)**. The initial design of BonFIRE utilized a point-scanning (PS) mode with tightly focused beams of both the MIR and the NIR lasers to ensure single-molecule detectability across the wide fingerprint and cell-silent regions (1300 cm^-1^ – 2400 cm^-1^) for double- and triple-bond vibrational modes (*23*). However, the serial-acquisition PS-BonFIRE is limited by its temporal resolution, making it challenging to capture rapid dynamic processes.

**Figure 1.**
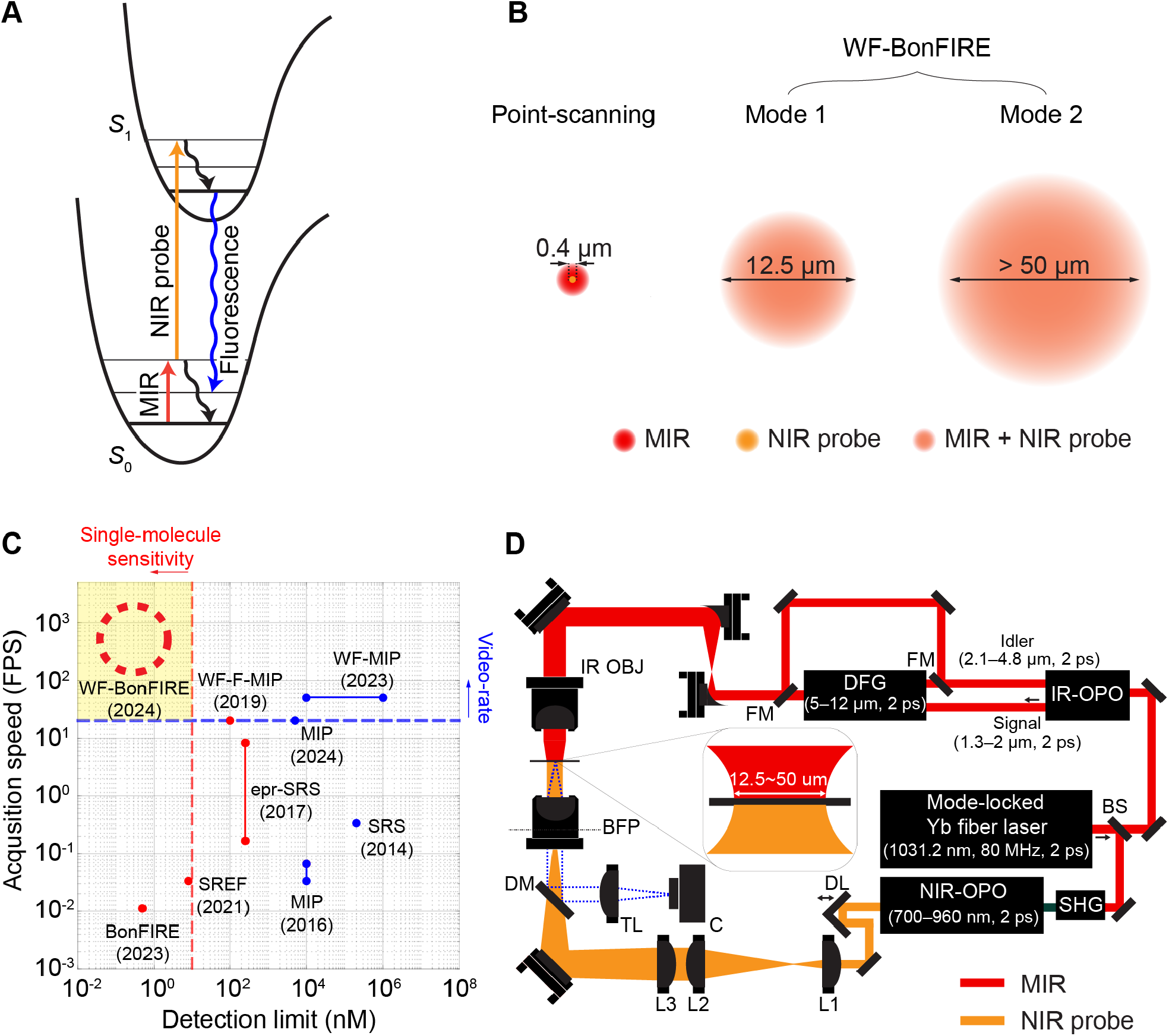
Principle and design of WF-BonFIRE. (A) Energy diagram of BonFIRE spectroscopy. S0 and S1 represent the ground and first electronic excited states. (B) Two implementation modes of WF-BonFIRE (Mode 1 & Mode 2), in comparison to the point-scanning mode. Not drawn to scale. (C) A scheme outlining the speed and sensitivity of WF-BonFIRE among other emerging bond-selective imaging modalities. Red dots indicate modalities that use exogenous dyes, while blue dots represent modalities that employ vibrational tags or are label-free. See Supplementary Table 1 for details. (D) WF-BonFIRE experimental scheme. Dotted blue line indicates fluorescence detection path. OBJ, objective; BFP, back focal plane; DM, dichroic mirror; TL, tube lens; C, camera; L, lens; DL, delay line; BS, beamsplitter; DFG, difference frequency generation; OPO, optical parametric oscillator; SHG, sum frequency generation; FM, flip mirror.

Here, we report wide-field (WF)-BonFIRE **(Fig. 1B)**. Through carefully balanced design and simulations of sensitivity against laser power, we push the imaging speed and the FOV limits of WF-BonFIRE to its maximum while achieving single molecule sensitivity, surpassing the performance of existing technologies (*20*–*22,25,26*) **(Fig. 1C)**. Such design facilitates up to 10000-fold faster frame acquisition compared to PS-BonFIRE. We demonstrate exceptional WF-BonFIRE imaging performance in cells, astrocytes, and in live neurons, capturing intricate structural details and networks with robust signal-to-noise ratios (SNRs). To further achieve kilohertz frame rate, we implement a temporal-delay modulation scheme that obtains up to 1500 frames per second (FPS) for WF-BonFIRE. We showcase the performance of temporal-delay modulation by tracking the Brownian motion of live Escherichia coli (E. coli). We anticipate that WF-BonFIRE will significantly push the boundaries of both chemical imaging and fluorescence imaging, facilitating high-speed and high-throughput imaging at the single-molecule level **(Fig. 1C)**.

## RESULTS

### Rational design and simulation of WF-BonFIRE

We first rationalize the feasibility of WF-BonFIRE especially in the single molecule regime. Contrary to conventional virtual-state mediated two-photon imaging techniques such as two-photon fluorescence and SRS microscopy, which face challenges in high-sensitivity WF implementation due to high photon-flux requirements, BonFIRE is a real vibrational state-mediated non-degenerate two-photon excitation process. Employing 2-ps laser excitation achieves a balance between bond-selectivity and efficient vibrational excitation, which competes with the picosecond vibrational relaxation lifetime **(Fig. S1)**. The up-conversion step is also highly efficient due to the large electronic absorption cross section. These efficient excitation steps hence alleviate the high photon flux requirement, thus allowing picosecond laser pulses to spread over a wider focal area. In addition, in PS-BonFIRE **(Fig. 1B, Point-scanning)**, the diffraction-limited MIR spot is significantly larger than that of the NIR probe laser, which results in more than 50% of MIR photons not being utilized for signal generation. Given the excess power of the NIR probe laser, the most straightforward design of WF-BonFIRE is to expand the NIR probe beam to match with the diffraction-limited spot of the MIR beam, thereby optimizing the utilization of MIR photons with opportunities for additional area expansion.

To quantitatively model the optimal FOVs, we next calculated the achievable signals and SNRs as a function of FOV using a high-sensitivity sCMOS camera **(Fig. S2, Eq S1-S3)**. At a 12.5 µm FOV, the achievable WF-BonFIRE signal from a single Rhodamine 800 (Rh800) molecule is estimated to reach 1 photon/ms with a SNR of 3, under a short camera exposure time of 25 ms. In situations where single-molecule sensitivity is not essential, the FOV can be further expanded. Hence, we investigated the maximum FOV for higher concentration samples. To this end, we introduced a pixel rate ratio, R (Eq. S4-S5), for comparing speeds between WF-BonFIRE and PS-BonFIRE. At the upper limit of R=1 where WF-BonFIRE theoretically matches the speed of PS-BonFIRE, FOV of WF-BonFIRE extends to 620 µm. However, reaching such large FOV results in reduced detection sensitivity, due to the inverse quadratic relationship between laser intensity and the illumination size for each beam **(Fig. S3)**.

Based on the above simulation, we implemented WF-BonFIRE in two modes, each providing optimal combination of sensitivity, speed and FOV for different biological applications. In Mode 1 **(Fig. 1B)**, we expanded and matched the sizes of the NIR probe and the MIR beams to a diameter of 12.5 µm for fast single-molecule imaging applications, paving the way toward WF-BonFIRE SMLM. In Mode 2 **(Fig. 1B)**, both beams are expanded and matched to 50 µm, sufficient to cover an entire mammalian cell while achieving an optimal balance between SNR and speed. At 50 µm FOV, the imaging speed of WF-BonFIRE is estimated to be over 150 times faster than PS-BonFIRE **(Fig. S3, R>150)**. When a larger FOV is needed, a parallel mosaicking approach is applied by precisely moving the piezo stage. Experimentally, we focused the probe beam onto the back focal plane of the objective to ensure uniform illumination **(Fig. 1D)**. Simultaneously, the IR beam underwent expansion using a lower numerical aperture (NA) MIR objective to align its spot size with that of the probe beam. System magnification was carefully determined to meet the requirements of the Nyquist theorem **(Fig. S4)**.

### Characterization of WF-BonFIRE (Mode 1) spectroscopy and imaging performance

We first validated WF-BonFIRE (Mode 1) by targeting the C=C bond (1598 cm^-1^) of Rh800, a NIR fluorescent molecule with an absorption peak at 696 nm **(Fig. S5)**. We tuned the NIR probe wavelength to 788 nm to achieve a sum frequency of NIR and MIR lasers that matches the absorption peak maximum **(Fig. S5)**. The WF-BonFIRE signal of Rh800-embedded spin-coated polymer film was generated by subtracting an image captured with a 20 ps temporal delay between MIR and NIR pulses from that with overlapping pulses (t_D_=0 ps) **(Fig. 2A & B)**. The background from temporally separated laser pulses is attributed to anti-Stokes fluorescence and photothermal signal, resulting from local temperature increases from the environment, both of which are constant across the temporal profile **(Fig. S6)** and exhibit no vibrational characteristics from the target molecules (*23*). The authenticity of the BonFIRE signal was further verified by the WF-BonFIRE spectrum, obtained by scanning the MIR laser wavelength, which closely aligns with the FTIR spectra from Rh800 in solution **(Fig. 2C)** featuring prominent peaks at 1505 cm^-1^ and 1598 cm^-1^, and by the linear power dependence on both MIR and NIR laser powers **(Fig. S7)**.

**Figure 2.**
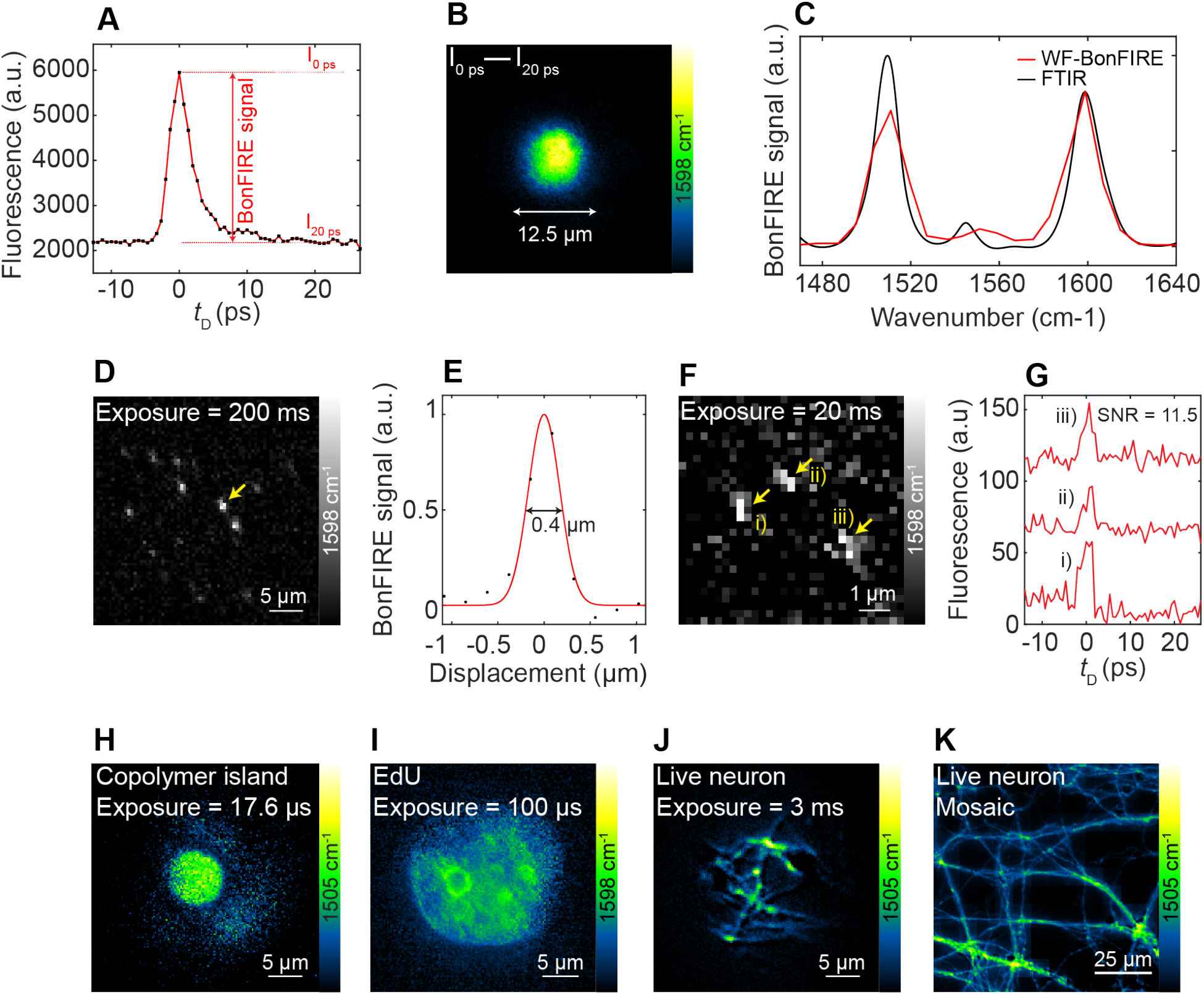
Characterization of WF-BonFIRE (Mode 1) spectroscopy and imaging performance. (A) WF-BonFIRE signal of Rh800 copolymer film as a function of temporal delay (tD). (B) WF-BonFIRE image of Rh800 copolymer film generated by subtracting two images acquired at temporal delays 0 ps and 20 ps (I0 ps and I20 ps shown in (A)). (C) Overlay of WF-BonFIRE (red) and FTIR (black) spectra of Rh800. (D) Single-molecule WF-BonFIRE image of Rh800 at 1598 cm^-1^. Exposure time: 200 ms. (E) Cross-section profile of a single molecule indicated in (D, yellow-arrowed). (F-G) Short-exposure (20 ms) WF-BonFIRE image (F) of single-molecule Rh800, with corresponding temporal profiles (G) for single molecules of i), ii), and iii) in (F). SNR = 11.5 (*N* = 3). (H) WF-BonFIRE images of Rh800 copolymer island at on-resonance (1505 cm^-1^). Exposure time: 17.6 μs. (I) WF-BonFIRE image targeting C=C vibration in ATTO680-click-labelled EdU in the nuclei of HeLa cells (I, 1,598 cm^−1^). (J-K) Single FOV (J) and mosaic (K) WF-BonFIRE images of Rh800-labelled mitochondria in live mouse neuronal cultures acquired at 1505 cm^-1^. Exposure time: 3 ms. Acquisition area in (K): 100 × 100 µm^2^.

To demonstrate the high sensitivity of WF-BonFIRE (Mode 1), single molecules of Rh800 were imaged with robust SNRs (> 48). This was achieved under an exposure time of 200 ms **(Fig. 2D, Fig. S8)** and even as short as 20 ms **(Fig. 2F & G)**, which is consistent with the predictions in Fig. S2. For single molecule measurements, the imaging speed of WF-BonFIRE exceeds that of PS-BonFIRE by over 200 times, underscoring its potential use in advanced super-resolution microscopy techniques like SMLM. By utilizing a single molecule as a point source, the spatial resolution of our WF-BonFIRE was characterized to be 400 nm **(Fig. 2E)**, confirming its diffraction-limited performance.

We then applied WF-BonFIRE (Mode 1) to rapidly image various samples ranging from copolymer films to live neurons. An exposure time of 17.6 µs was achieved when imaging a Rh800-labeled copolymer film, consisting of phase-separated polystyrene and polymethylmethacrylate segments **(Fig. 2H)** with high bond-selectivity **(Fig. S9A)**. The imaging speed was 10,000 times faster than PS-BonFIRE, closely aligning well with our earlier predictions **(Fig. S3)**. In these experiments, a single polymer island was captured, which was approximately the size of small mammalian cells (∼10 μm). Additionally, we imaged a single nucleus of a HeLa cell using microsecond-level exposure time **(Fig. 2I & Fig. S9B, 100 μs)**, using ATTO680 click-labeled 5-ethynyl-2’-deoxyuridine (EdU) in newly synthesized DNA. The short exposure time is expected to enable biological imaging at kilohertz speed, a feat challenging for existing chemical-selective imaging modalities. Leveraging its high sensitivity and speed, WF-BonFIRE further demonstrated live neuron imaging at an exposure time of 3 ms, revealing distributions of mitochondria labeled with Rh800 **(Fig. 2J & K, & Fig. S9C)**. Such speeds fulfill the necessary requirements for resolving fast dynamics in cells, such as voltage imaging in neurons. Additionally, by mosaicking single FOV WF-BonFIRE images, a network of live neurons was captured within ∼30 seconds, enabling extensive imaging of a large area (100 × 100 µm^2^) of live cells **(Fig. 2K)**.

### Large FOV WF-BonFIRE imaging (Mode 2)

We then explored the expanded FOV capabilities of WF-BonFIRE (Mode 2), which more effectively captures a larger area for biological samples. Imaging of a Rh800 copolymer sample is achieved at an exposure time of 500 μs **(Fig. 3A)**. Additionally, a 200 × 200 µm^2^ area was acquired within 1.3 s using mosaicking **(Fig. 3B)**. Although Mode 2 lacks single-molecule sensitivity, it effectively imaged a wide variety of biological targets, including low-abundance proteins at micromolar concentrations or lower, within an expanded FOV **(Fig. 3C-H)**. Using ATTO680-immuno-labeled GFAP, a key marker protein for astrocytes, we obtained an exposure time of 100 ms for single FOV WF-BonFIRE (Mode 2) imaging that targeted the 1598 cm^-1^ of the C=C vibration **(Fig. 3C & D)**. This enabled capturing a 200 × 200 um^2^ area within 10 s **(Fig. 3D, Mosaic)**, with high bond-selectivity **(Fig. 3D, inset)**. Similarly, ATTO680-labeled α-tubulin **(Fig. 3E & F)** and EdU **(Fig. 3G & H)** were imaged with WF-BonFIRE (Mode 2). The high SNR enabled resolving clear tubulin **(Fig. 3E)** and nuclei **(Fig. 3G)** structures, providing detailed insights into their spatial organization. Here, a single FOV image successfully captured an entire HeLa cell **(Fig. 3E)** and multiple nuclei **(Fig. 3G)**.

**Figure 3.**
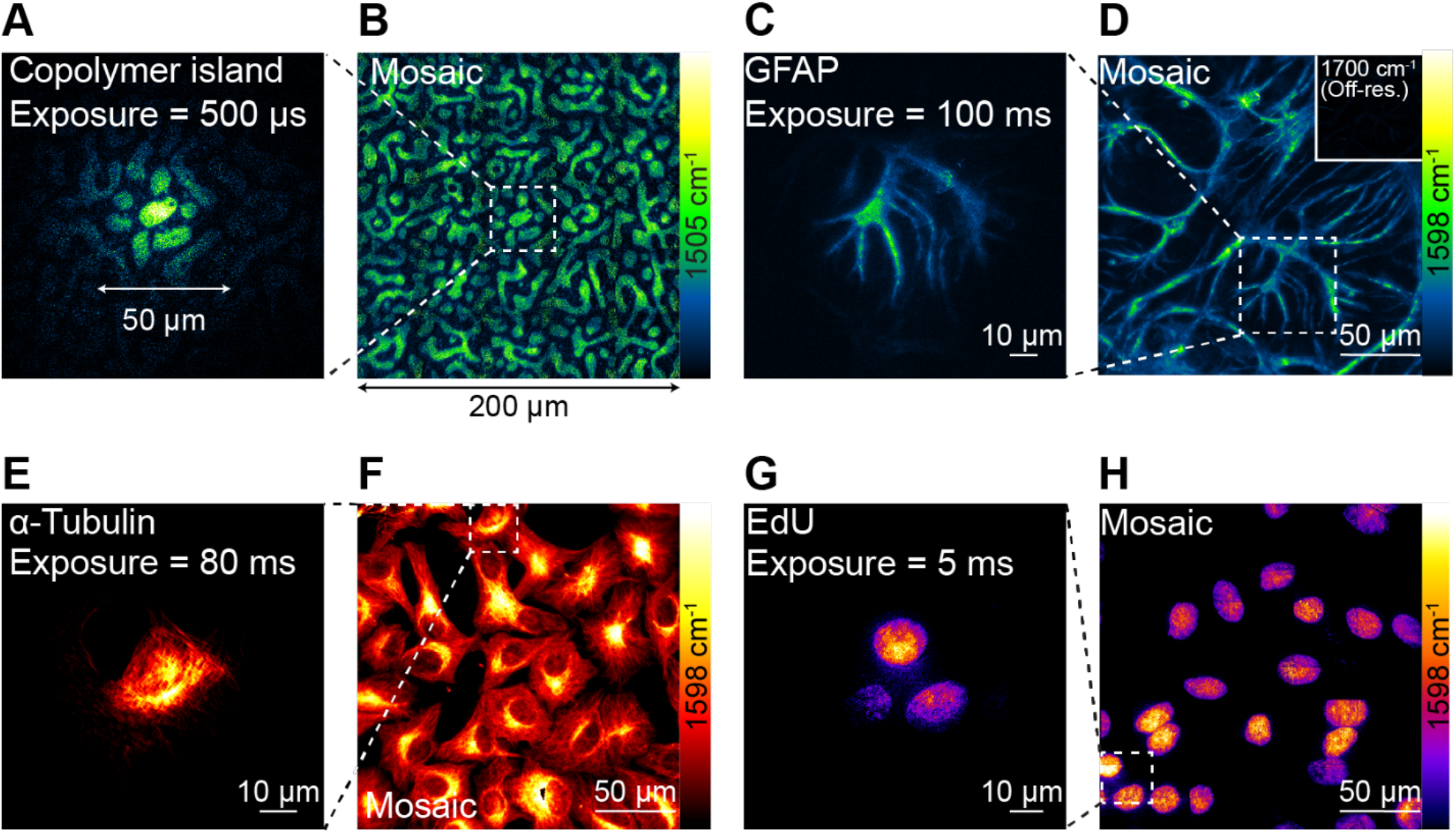
Large FOV WF-BonFIRE imaging (Mode 2). (A-B) Single FOV (A) and mosaic (B) WF-BonFIRE images of Rh800 copolymer film acquired at 1505 cm^-1^. Exposure time: 500 µs. Acquisition area in (B): 200 × 200 µm^2^. (C-H) Single FOV (C, E, G) and mosaic (D, F, H) WF-BonFIRE images targeting C=C vibration (1598 cm^-1^) for ATTO680-immunolabeled GFAP in mouse neuronal co-cultures (C & D. Exposure time: 100 ms. Acquisition area in (D): 200 × 200 µm^2^); ATTO680-immunolabelled α-tubulin in HeLa cells (E & F. Exposure time: 80 ms. Average: 5 frames. Acquisition area in (F): 200 × 200 µm^2^); and ATTO680-click-labelled EdU in the nuclei of HeLa cells (G & H. Exposure time: 5 ms. Average: 16 frames. Acquisition area in H: 200 × 200 µm^2^).

### Kilohertz WF-BonFIRE imaging with newly-developed temporal-delay modulation

A major advantage of WF microscopy is its ability to perform live-cell dynamic imaging at kilohertz frame rates surpassing video-rate FPS, which is valuable for applications such as tracking bacterial movement or mapping the neuronal action potentials. Although WF-BonFIRE achieved microsecond acquisition speed for a single image, a significant bottleneck to reaching kilohertz framerate is the requirement to subtract subsequent images between the temporal on (t_D_=0 ps) and off states (t_D_=20 ps) to effectively remove the flat photothermal background, which accumulates over a slower timescale (*23*). The adjustment of the temporal delay involves tuning a delay stage, which requires an additional 0.2-second mechanical settling **(Fig. S10)**, thereby hindering the achievement of high-speed dynamic imaging.

To address this challenge, we developed a temporal-delay modulation scheme that allows instantaneous modulation of pulse delays without relying on the physical position of the mechanical delay line **(Fig. 4A & B)**. To achieve this, the NIR probe beam was split into two pulse trains of equal intensity using a beam splitter **(Fig. 4A & B, BS1)**. The equal intensity for each beam arm was carefully calibrated using a continuously variable neutral density (ND) filter. One path was deliberately lengthened by Δl relative to the other **(Fig. 4B)**, introducing an additional delay (tD) of 20 ps compared to the pulse trains from the shorter path with t_D_=0 ps **(Fig. 4A)**. To synchronize the imaging, a chopper rotating at half the frequency of camera frame rate was employed to alternately allow pulse trains from only one path to reach the sample at a time, while blocking the other **(Fig. 4A & B, Chopper)**. These two beams were then recombined using a second beam splitter **(Fig. 4A & B, BS2)** and directed towards the sample, producing pulse trains with alternating temporal delays **(Fig. 4C, Recombined at BS2)**. The chopper was precisely synchronized with the camera, ensuring that one image was captured with only the first pulse train at t_D_ = 0 ps, followed by another image with the second pulse train at t_D_ = 20 ps, **(Fig. 4D & E)**.

**Figure 4.**
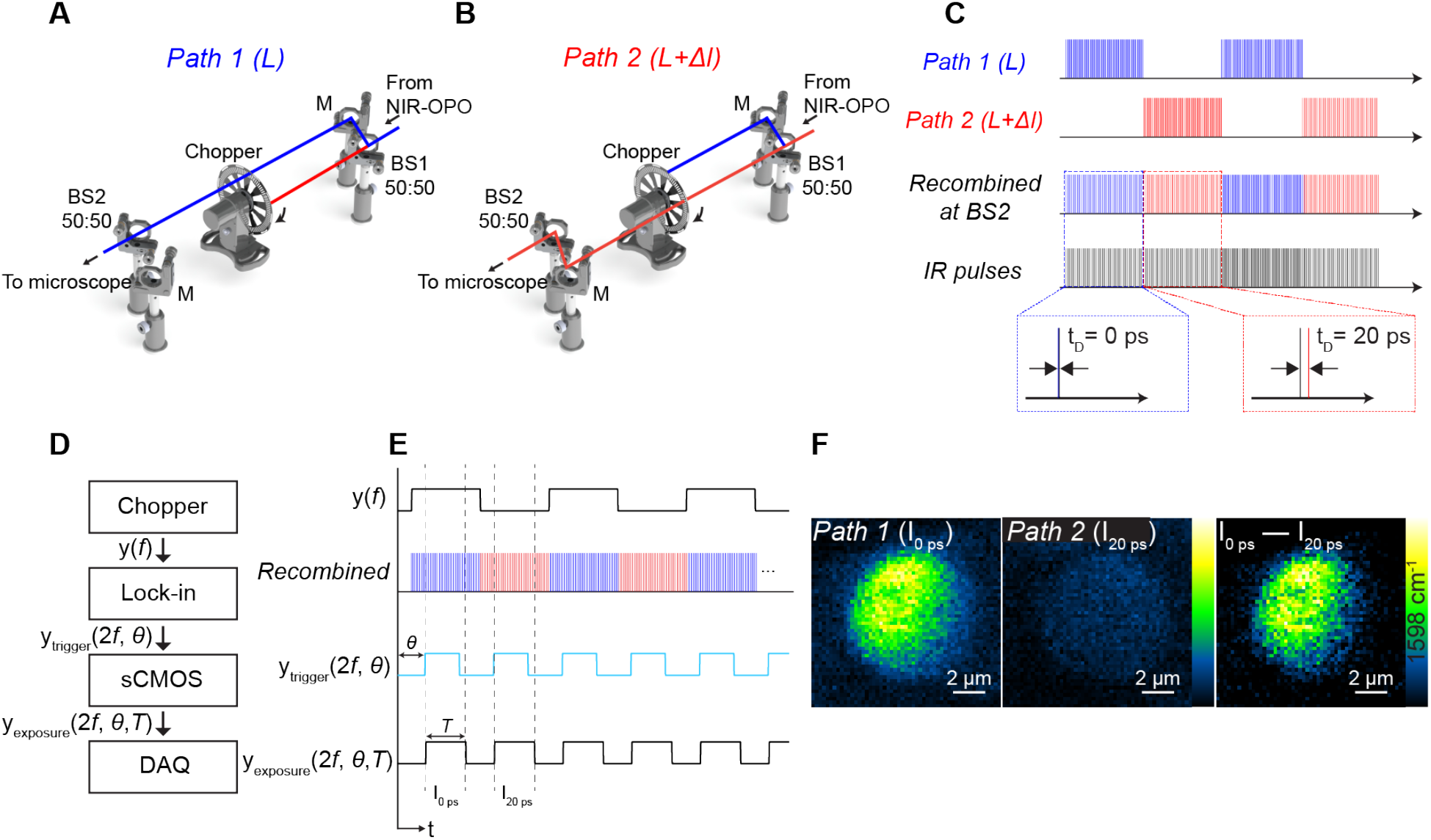
Kilohertz WF-BonFIRE imaging using temporal delay modulation. (A-C) Experimental set-up of temporal delay modulation scheme. NIR laser pulses, divided into Path 1 (A, blue) and Path 2 (B, red), which have the exact same intensity but with different path lengths (Δl), introduce different temporal delays (0 ps and 20 ps) relative to the pulse trains of the MIR pulses after recombination at BS2 (C). A chopper is precisely aligned and synchronized to ensure that only one path of NIR pulse trains reach the sample at any given time. BS: Beam splitter. (D) Wiring diagram for camera-chopper synchronization. (E) Synchronization timing chart. (F) Kilohertz (FPS, 1500 Hz) WF-BonFIRE image (I_0 ps_ – I_20 ps_) acquired using temporal delay modulation, through subtraction of subsequent camera frame images, where one image is from Path 1 pulse trains (Path 1 (I_0 ps_)) and the other image is from Path 2 pulse trains (Path 2 (I_20 ps_)).

In Figure 4D, we present a wiring diagram for control signals that govern camera and lock-in synchronization within the system. The reference output from the chopper (y(*f*)) is doubled in frequency by a lock-in amplifier, and this digital output (y_trigger_(2*f, θ*)) serves as the trigger for the sCMOS camera’s exposure **(Fig. 4D & E, y_exposure_(2*f, θ, T*)**). The critical elements for achieving precise synchronization are the camera’s exposure timing output and the recombined alternating pulse train **(Fig. 4E)**. To ensure seamless coordination, adjustments are made to the lock-in phase (*θ*) to account for the chopper’s rise and fall times, during which the beam partially obstructs. Moreover, the exposure duration (*T*) is set to approximately 60% of the reciprocal of the doubled chopper frequency (1/2*f*).

Utilizing the temporal delay modulation scheme, we performed real-time WF-BonFIRE (Mode 1) imaging on a Rh800 copolymer sample **(Fig. 4F)**. Subsequent camera frames corresponding to Path 1 (I_0 ps_) and Path 2 (I_20 ps_) delays were captured according to the chopper’s modulation frequency **(Fig. 4E)**. Consequently, a background-free WF-BonFIRE image **(Fig. 4F, I_0 ps_-I_20 ps_)** was directly obtained through subtraction. As a control, dark images were consistently confirmed by setting the time delays of Path 1 and Path 2 to the temporal-off position, indicating the absence of artifacts or power difference between the two paths **(Fig. S11)**. Using the temporal delay modulation, we achieved kilohertz real-time WF-BonFIRE imaging at 1500 FPS **(Fig. 4F)**. To demonstrate WF-BonFIRE’s dynamic imaging capabilities beyond video rate, we applied it to track the random Brownian motion of Rh800-stained E. coli —a critical aspect of microbial behavior in liquid environments. The diffusion coefficient of E. coli, typically determined by the Stokes-Einstein equation, falls within the range of 10 to 100 µm^2^/s (*27*). To effectively capture and analyze this rapid microscopic motion, high-speed microscopy with frame rates exceeding several tens of frames per second is required. We acquired WF-BonFIRE images at 150 FPS. In Fig. 5, eight consecutive frames within 50 milliseconds are displayed, confirming that WF-BonFIRE can precisely track the rapid movement of E. coli without motion artifacts **(Fig. 5, Movie S1)**.

**Figure 5.**
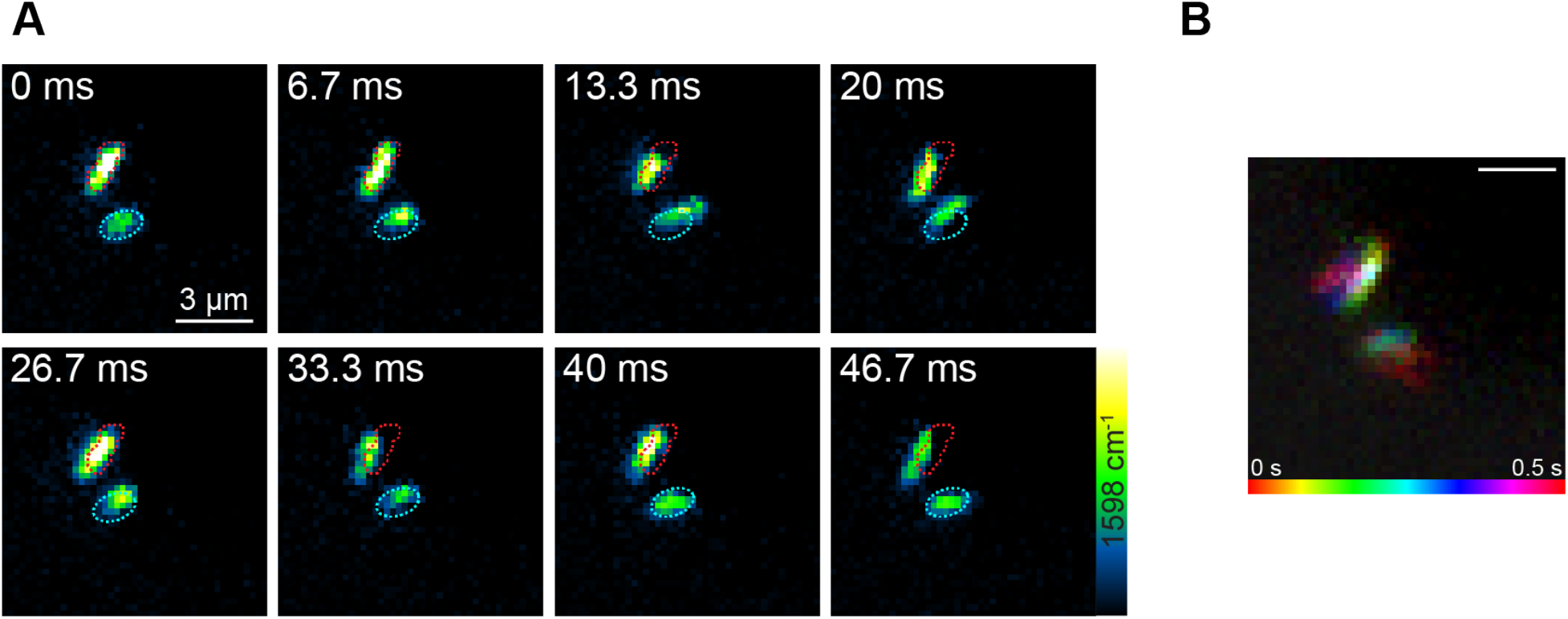
WF-BonFIRE imaging capturing Brownian motion in Rh800-stained E. coli at 150 frames per second (FPS) using temporal delay modulation. (A) WF-BonFIRE images at sequential time points. Red and blue dotted profiles indicate positions of the E. coli at t = 0 ms frame. (B) Temporal color-coded image showing the movement of E. Coli over a period of t = 0 to 0.5 s. Scale bar, 3 µm.

## DISCUSSION

In this manuscript, we present wide-field bond-selective fluorescence imaging that achieves high sensitivity and speed. This advancement is attributed to the efficient vibronic transitions of WF-BonFIRE, which reduce the typically high photon flux requirements of multiphoton microscopy. We demonstrate significant increases in imaging speed using WF-BonFIRE compared to PS-BonFIRE, for both polymer films at high concentrations and single-molecule samples. Key results include achieving a 20-millisecond exposure time for single-molecule WF-BonFIRE imaging with robust SNRs, and rapid WF-BonFIRE acquisition to capture fine structures in live neurons and cells at low concentrations. This method achieved large FOV imaging (200×200 μm^2^) within several seconds, exceeding point-scanning capabilities. Moreover, by introducing a temporal delay modulation scheme, WF-BonFIRE achieved imaging kilohertz speed up to 1500 FPS, far exceeding video rates. This development enables the tracking of Brownian motion in E. coli, which requires millisecond-level temporal resolution, and offers potential to capture other rapid dynamic processes such as neuronal firing.

Imaging speed is influenced by the SNR, which is primarily determined by the photon flux of MIR and NIR probe lasers where shot noise is the limiting factor. In scenarios where detector noise-limited conditions prevail and the available laser power exceeds necessity, increasing the FOV by a factor of M (in one dimension) improves imaging speed by M^2^. However, our system primarily operates in a source-limited regime where an increased FOV decreases photon flux and signal, resulting in a net decrease in overall imaging speed despite gains from spatial multiplexing. For this reason, we have implemented two distinct operational modes to accommodate various application needs. Mode 1 is utilized for situations that demand high photon flux, including single-molecule imaging and fast dynamic imaging. Mode 2 is selected for simultaneous detection across larger areas, suited for relatively more abundant molecular targets. Additionally, using a larger FOV reduces the number of stage steps needed to cover a large acquisition area (e.g. 200×200 μm^2^). As an outlook, adopting a higher-power laser could transition our system to a detector-limited regime, enabling larger FOVs beyond 50 μm.

Although the imaging speeds of up to 1500 FPS have been achieved, the full potential of this technique has not yet been realized due to limitations of the current camera technology and frequency constraints of chopper rotation. For instance, the sCMOS camera used in our experiments is limited to a maximum speed of 2100 FPS with external triggering. Achieving speeds greater than 10000 FPS presents challenges with the current temporal delay modulation scheme, as the chopper’s maximum speed is limited to 10 kHz. Nonetheless, employing electro-optic modulators (EOM) and quartz crystals could provide a solution by enabling temporal delay modulation at significantly higher frequencies, potentially overcoming these limitations (*28*).

An important future application of WF-BonFIRE lies in super-resolution bond-selective imaging. In chemical imaging, achieving sub^-1^00 nm spatial resolution has been challenging due to limited SNR (*29*–*32*). Combining chemical imaging with techniques such as STED and RESOLFT often results in photobleaching and limits multiplexing capabilities. In contrast, the sensitivity and speed of WF-BonFIRE offer considerable opportunities for super-resolution bond-selective imaging, especially using single-molecule localization microscopy (SMLM), which could substantially minimize photobleaching effects (*33*). When combined with SMLM, WF-BonFIRE could enable highly multiplexed super-resolution microscopy with local environment sensing (*24*), addressing a challenge for current fluorescence techniques.

## MATERIALS AND METHODS

### Materials

ATTO dyes were purchased from ATTO-TEC. Rhodamine 800 was purchased from Sigma Aldrich. All dyes were aliquoted in DMSO as stock solutions (10 mM) upon receipt and stored at −20 °C. The following primary antibodies were used: anti-α-tubulin in rabbit (ab18251, Abcam) and anti-GFAP in mouse (3670S, Cell Signaling Technology). The following secondary antibodies were used: goat anti-mouse antibody (31160, Invitrogen) and goat anti-rabbit antibody (31210, Invitrogen).

### Experimental setup of WF-BonFIRE

The laser sources for WF-BonFIRE are identical to that of PS-BonFIRE^1^. A 1.8 ps, 80 MHz, 1031.2 nm mode-locked Yb fiber laser (aeroPULSE PS10, NKT Photonics, Copenhagen, Denmark) was used as a seed laser for both NIR and MIR optical parametric oscillators (OPOs), providing wide wavelength turnabilities. The frequency doubled beam was used to pump the NIR-OPO (picoEmerald, Applied Physics and Electronics), which tunes from 700-960 nm. IR-OPO (Levante IR, Applied Physics and Electronics, Berlin, Germany) generates an idler beam that tunes from 2.1-4.8 µm (2083-4762 cm^-1^). Differential frequency generation (HarmoniXX DFG, Applied Physics and Electronics) inputs signal and idler beams of the IR-OPO to generate MIR wavelengths from 5^-1^2 µm (833-2000 cm^-1^).

For wide-field illumination, the NIR beam was focused onto the back focal plane of the NIR objective (XLPLN25XWMP2, Olympus) using plano-convex lens to ensure homogeneous illumination. The location of the back focal plane was confirmed by monitoring the collimation of the beam after the objective at varying lens position. A 2000 mm (LA1258-B-ML, Thorlabs) and 500 mm (LA1908-B-ML, Thorlabs) focal length lens were used to achieve 12.5-µm (Mode 1) or 50-µm (Mode 2) FOVs, respectively. The MIR beam was loosely focused onto the sample plane using a 6.35 mm (39-469, Edmund optics) or a 20 mm (LA7733, Thorlabs) focal length lens to achieve 12.5-µm (Mode 1) and 50-µm (Mode 2) FOVs, respectively. For the imaging path, a sCMOS camera (C15440-20UP, Hamamatsu Photonics) was installed at the focal plane of the tube lens with focal length of 200 mm. The magnification of the imaging system was 200 mm/7.2 mm = 27.78x, confirmed by measuring the effective pixel size. Fluorescence was separated from excitation using a dichroic mirror (FF738-FDi01, AVR optics) and bandpass filter (FF01-665/150, AVR optics). A custom LabVIEW program was used for all data acquisition, including stage scan, delay line movement, and camera capture.

For temporal delay modulation, a pair of pellicle beamsplitters (BP145B2, Thorlabs) were used to divide the beam into two paths. To ensure equal power in both beams, a continuously variable neutral density (ND) filter (NDL^-1^0C-2, Thorlabs) was placed in one of the paths and adjusted until the fluorescence signals measured using a photomultiplier tube (PMT1002, Thorlabs) from both paths were identical. The XY translation stage of the chopper (MC2000B, Thorlabs) was then fine-tuned to make both beams equidistant from the chopper’s center. The optimal position of the chopper was found by minimizing the demodulated anti-stokes fluorescence signal at the chopper frequency. The reference output from the chopper was frequency-doubled using a lock-in amplifier (HF2LI, 50-MHz bandwidth, Zurich), which was then used to trigger the sCMOS camera. To achieve perfect synchronization between the chopper and the camera, the phase of the lock-in output was adjusted. This setup ensured that only the pulse trains from a single path reached the sample at any given time.

### Preparation of single-molecule and polymer samples

For single molecule samples, PVA (363138, Sigma) solution of 2.7 mg/mL in H2O was used to dilute Rh800 (83701, Sigma) solution in DMSO to 1 pM. Rh800-PVA solution was spin coated onto a CaF2 window (CAFP10-0.35, Crystran) at 5000 RPM for 30 seconds. For copolymer film sample, 12 mg/mL polystyrene (430102, Sigma) and 18 mg/mL PMMA (200336, Sigma) in toluene were mixed. After dissolving Rh800 with the mixture to reach a final concentration of 100 µM, Rh800-PS-PMMA solution was spin coated (BSC^-1^00, MicroNano Tools) onto a CaF2 window at 1200 RPM for 30 seconds.

### Preparation of biological samples

HeLa-CCL2 (ATCC) cells were seeded onto 10 mm diameter 0.35-mm-thick CaF2 windows at a cell density of 10^5^ cells/mL and were cultured in Dulbecco’s modified Eagle medium (DMEM) at 37 °C with 5% CO_2_. The DMEM mixture was composed of 90% DMEM (11965, Invitrogen), 10% fetal bovine serum (FBS; 10082, Invitrogen), and 1× penicillin/streptomycin (15140, Invitrogen). For HeLa cells with EdU labeling, the medium was switched to FBS-free DMEM (Gibco) for 20–22 hours to synchronize the cell cycle. After synchronization, the medium was reverted to the original DMEM and EdU (10 mM stock in H2O) was added at a concentration of 200 μM for 20–24 hours. The cells were then fixed with 4% paraformaldehyde (PFA) for 20 min, and the PFA was removed using Dulbecco’s phosphate-buffered saline (DPBS).

### Neuron culture

Primary hippocampal neurons were isolated from neonatal Sprague–Dawley rat pups using a Caltech-approved protocol (IA22^-1^835) by the Institutional Animal Care and Use Committee (IACUC). The brains were removed and immersed in ice-chilled Hanks’ balanced salt solution (Gibco) in a 10-cm Petri dish. Under a dissection microscope, the hippocampi were separated, finely minced to approximately 0.5 mm pieces, and digested in 5 mL of 0.25% Trypsin-EDTA (Gibco) at 37°C in a 5% CO2 incubator for 15 min. After aspiration of the Trypsin-EDTA, the tissue was quickly neutralized with 2 mL of DMEM containing 10% FBS. The tissue pieces were then gently moved into 2 mL of neuronal culture medium (Neurobasal A, B-27 and GlutaMAX supplements, Thermo Fisher, with 1× penicillin-streptomycin) to dissociate the cells. The resultant cell suspension was further diluted to a final density of 9 × 10^4^ cells/mL using the same medium. Each well of a 24-well plate, containing pre-coated CaF_2_ windows, received 0.7 mL of this suspension. The CaF2 windows had been prepared by incubating them with 100 μg/mL poly-d-lysine (Sigma) at 37°C and 5% CO_2_ for 24 hours, followed by a laminin mouse protein (Gibco) layer at 10 μg/mL, also at 37°C and 5% CO_2_ overnight. After rinsing twice with ddH2O and drying in a biosafety cabinet at room temperature, the neurons were maintained with a half-medium exchange every four days. At day 14 in vitro (DIV14), neurons were fixed with 4% paraformaldehyde (PFA) for 20 min, washed with DPBS, and could be stored in DPBS at 4 °C for several days.

### Click reaction labeling of EdU incorporated cells

Initially, the EdU-labeled cells underwent permeabilization using 0.2% X^-1^00 (T8787, Sigma) for 20 min. They were then washed with 2% BSA in PBS. Subsequently, the cells were incubated in a reaction buffer containing 10 µM of ATTO680, prepared as specified in the click reaction buffer kit (Thermo Fisher, C10269). After a 30-minute incubation at room temperature, the cells were washed again with 2% BSA in PBS prior to imaging.

### Immunolabeling of fixed HeLa and neurons

Fixed cells were first permeabilized using 0.1% Triton X^-1^00 (T8787, Sigma) for 20 min. After blocking for 1–3 h in 10% goat serum/1% BSA/0.3 M glycine/0.1% PBST, the cells were incubated overnight at 4 °C in 10 μg mL−1 primary antibody in 3% BSA. After washing with PBS, the cells were blocked using 10% goat serum in 0.1% PBST for 1–3 h, followed by overnight incubation at 4 °C in ∼10 μg mL−1 secondary antibody in 10% goat serum. The cells were blocked with 10% goat serum for 30 min and dried before imaging.

