## Supplementary Material for "Wide-field bond-selective fluorescence imaging: from single-molecule to cellular imaging beyond video-rate"

Lee, Dongkwan *et al.*

**This PDF file includes:**

Table S1
Figs. S1 to S11
Caption of Movie S1

**Other Supplementary Materials for this manuscript include the following:**
Movie S1

### Table S1. Summary of sensitivity and speed of emerging bond-selective imaging modalities

| Table of Reported Imaging Speeds* | | | | | |
|---|---|---|---|---|---|
| Reference (Year) | Source | Pixel dwell time (µs) | X dimension | Y dimension | FPS |
| MIP (*22*) (2023) | Fig. 3A | 2 | 150 pixels | 100 pixels | 20 |
| FM-SREF (*29*) (2021) | Fig. 5a | 1000 | - | - | 0.033 |
| BonFIRE (*23*) (2023) | Fig. 5c | 3000 | - | - | 0.011 |
| epr-SRS (*13*) (2017) | Fig. 1c | 4 | - | - | 8.3 |
| | Fig. 2c | 200 | - | - | 0.17 |
| SRS (*34*) (2014) | All imaging | 100 | - | - | 0.33 |
| MIP (*19*) (2016) | Fig. 3A | 1000 | - | - | 0.033 |
| | Fig. 4C | 500 | - | - | 0.067 |
| | Fig. 5A | 500 | - | - | 0.067 |
| WF-F-MIP (*35*) (2019) | Fig. 5b | - | 70 µm | 40 µm | 20 |
| WF-MIP (*21*) (2023) | Fig. 5b | - | 87 µm | 87 µm | 50 |
| WISE (*25*) (2024) | Fig. 5F | - | 30 µm | 30 µm | 0.1 - 2 |
| WF-BonFIRE (2024) | Fig. 5 | - | 12.5 µm | 12.5 µm | 150 |

* The imaging speeds listed refer exclusively to imaging of biological samples. When the FPS was not explicitly stated for the point-scanning system, it was calculated based on 150 × 100 pixel dimensions to ensure a fair comparison.

| Table of Reported Sensitivity | | |
|---|---|---|
| Reference (Year) | Source | Detection limit (nM) |
| MIP (*36*) (2024) | Fig. 2e, 2f | 5000 |
| SREF (*14*) (2019) | Fig. 2g | 8 |
| BonFIRE (*23*) (2023) | Fig. 2h | 0.5 |
| epr-SRS (*13*) (2017) | Fig. 1b | 250 |
| SRS (*33*) (2014) | Fig. 1b | 200,000 |
| MIP (*19*) (2016) | Fig. 2B | 10,000 |
| WF-F-MIP (*35*) (2019) | Estimated | 100 |
| WF-MIP (21) (2023) | Estimated | 10,000 – 1,000,000 |
| WISE (*25*) (2024) | Not reported, hence not included in Fig. 1c | |
| WF-BonFIRE (2024) | Fig. 2D & F | Single-molecule level |

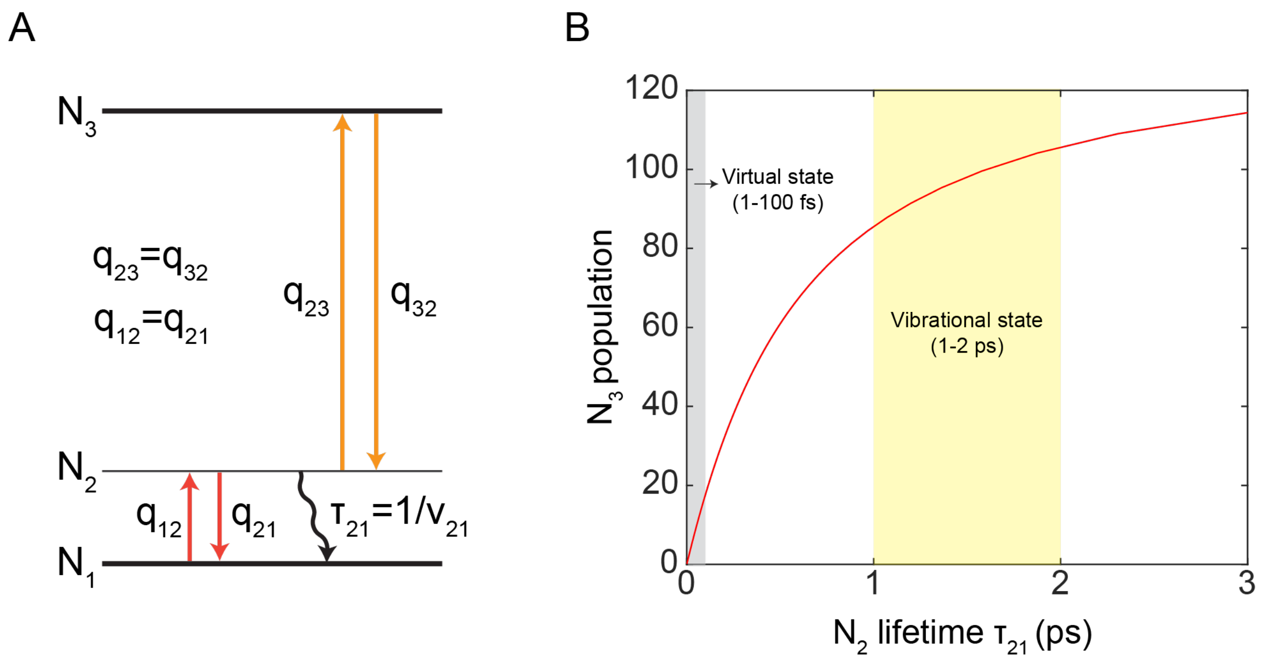


**Figure S1. BonFIRE signal calculation with varying lifetimes of intermediate state**. (A) The energy diagram of a three-level system used for the simulation. $N_1$ and $N_3$ indicate the ground and excited electronic state; $N_2$ indicates the intermediate state, which is either a virtual state or vibrational state in our analysis. (B) $N_3$ state population as a function of vibrational lifetime of $N_2$ state ($\tau_{21}$) calculated from the three-level rate equation simulations (*23*). On-sample laser powers: NIR Probe=0.1 mW and MIR=10 mW. The pulse duration is 2 ps for both lasers. Grey and yellow highlights the relevant temporal regions for typical lifetimes of virtual states (1-100 fs, gray) and vibrational states (1-2 ps, e.g., double bonds and nitriles, yellow). Significant increase in $N_3$ population is observed as intermediate state lifetimes shifts from virtual (gray) to vibrational state (yellow), indicating efficient vibronic excitation.

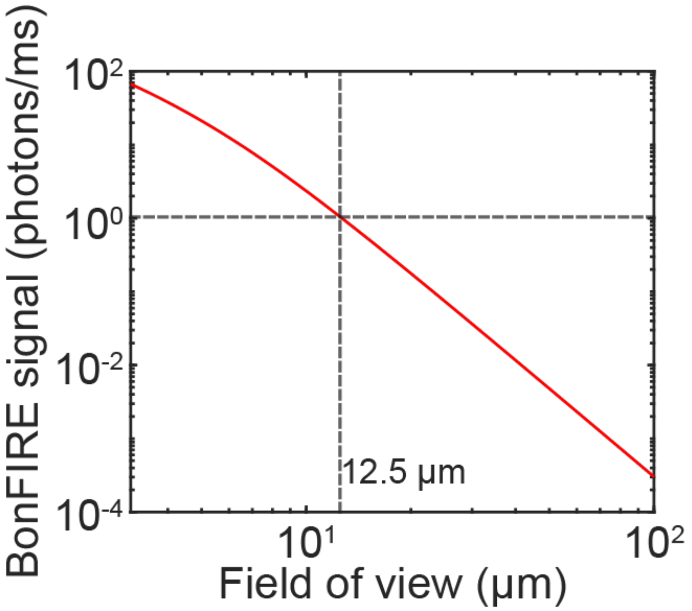


**Figure S2**. **WF-BonFIRE signal estimation at varying field-of-view (FOV) size using Rh800.** Three-level rate equations were used to calculate the BonFIRE signal (photons/ms) (*23*). Maximum on-sample MIR (10 mW) and NIR probe powers (150 mW) were used for the simulation. The pulse duration is 2 ps. At FOV of 12.5 μm, BonFIRE signal was computed to be 1 photon/ms. Signal-to-noise ratio (SNR) can be computed by the following equations.

$$SNR = \frac{QE \cdot N \cdot t}{\sqrt{\sigma_{shot\ noise}^2 + \sigma_{read\ noise}^2 + \sigma_{dark\ noise}^2}} \tag{S1}$$

$$\sigma_{shot\ noise} = \sqrt{QE \cdot N \cdot t} \tag{S2}$$

$$\sigma_{dark\ noise} = \sqrt{I_d \cdot t} \tag{S3}$$

where $QE$ is quantum efficiency, $N$ is number of photons per pixel per millisecond, $t$ is exposure time, and $I_d$ is dark current. At 12.5 μm FOV, SNR of 3 can be achieved with exposure times of 25 ms, indicating single-molecule sensitivity while achieving wide-field detection.

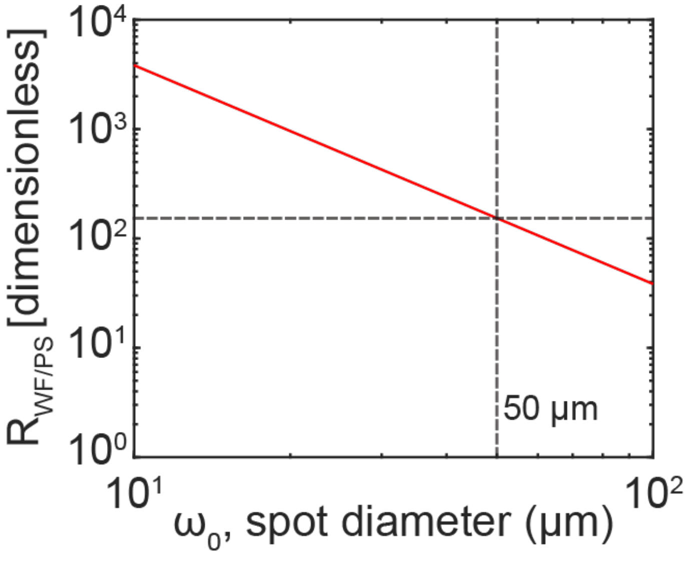


**Figure S3**. **WF-BonFIRE speed estimation relative to PS-BonFIRE as a function of FOV.** FOV is indicated as the spot diameter of the laser beams, $\omega_0$. Due to the quadratic reduction in photon flux across both probe and IR beams, WF-BonFIRE signal is expected to diminish quickly with increasing FOV. Although this would make single molecule detection infeasible, it should offer superior speed than the PS-BonFIRE for bio-imaging applications that do not require single-molecule sensitivity. Theoretically, the upper limit of FOV for WF-BonFIRE is when the imaging acquisition speed of WF-BonFIRE becomes equal (R=1) and then slower comparing to PS-BonFIRE. To perform the comparison, we followed a previous report that defined a pixel rate and pixel rate ratio $R$ as following (*37*):

$$pixel\ rate\ \equiv (\ total\ acquisition\ time\ normalized\ by\ number\ of\ pixels\ and\ SNR^2)^{-1} \qquad \text{(S4)}$$

$$R_{WF/PT} \equiv \frac{pixel\ rate\ for\ widefield}{pixel\ rate\ for\ point\ scanning} = \frac{\left(\frac{t_{WF}}{n_{WF}\cdot SNR_{WF}^2}\right)^{-1}}{\left(\frac{t_{PS}}{n_{PS}\cdot SNR_{PS}^2}\right)^{-1}} = \frac{n_{WF}}{n_{PS}}\frac{t_{PS}}{t_{WF}}\left(\frac{SNR_{WF}}{SNR_{PS}}\right)^2 \qquad \text{(S5)}$$

where $n$ is the number of pixels, $t_{PS}$ is the pixel dwell time, $t_{WF}$ is the camera exposure time, and $SNR$ is the signal-to-noise ratio. Assuming the system shot noise is the dominant noise source, Equation (S1) is simplified to:

$$SNR\ = \sqrt{QE\cdot C\cdot (k\cdot \sigma\cdot I_{IR}\cdot I_{probe})\cdot t} \qquad \text{(S6)}$$

where $QE$ is quantum efficiency, $C$ is concentration, $k$ is a scaling constant, $\sigma$ is the BonFIRE cross-section, $I$ is the peak intensity, and $t$ is the acquisition time. Combining Equation (S5) and (S6),

$$R = \frac{16}{\pi} \cdot \frac{A_{IR_{PS}} \cdot P_r}{\omega_0^2} \; where \; P_r \equiv \frac{P_{probe\_WF}}{P_{probe\_PS}} \qquad \text{(S7)}$$

where $A_{IR\_PS}$ is diffraction-limited area at MIR wavelength, $\omega_0$ is spot diameter, and $P_r$ is the ratio of the probe power used for wide-field and point-scanning. $P_r$ is determined by various factors (e.g. SNR, concentration of the sample, saturation) and experimentally determined to be ~5000 for many biological samples. Using Equation (S7), $R$ is calculated to be 153 at $\omega_0 = 50$ µm and $P_r$ = 5000, achieving more than two orders of magnitude increase in imaging speed compared to point-scanning. In theory, the FOV reaches 620 µm when R = 1, at which wide-field is the same speed compared to point-scanning. However, such large FOV would significantly sacrifice the detection sensitivity as reasoned above. We hence choose to perform Mode 2 WF-BonFIRE at $\omega_0 = 50$ µm to target both high sensitivity and fast speed.

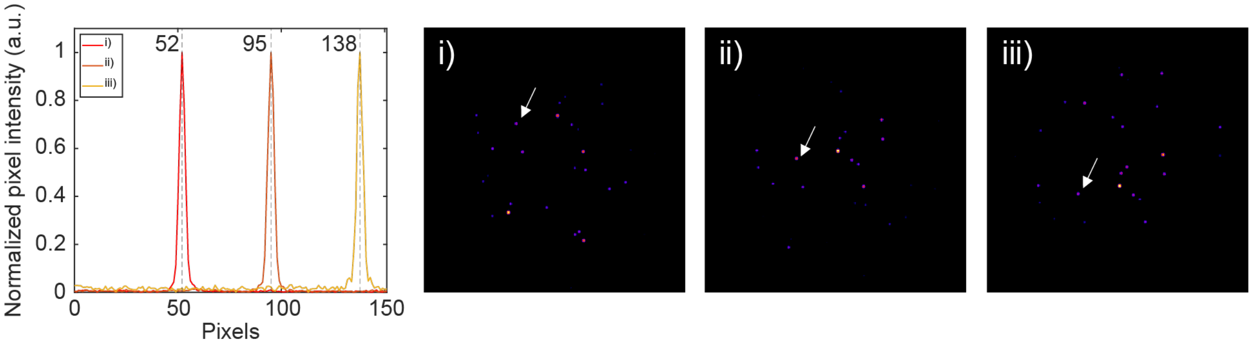


**Figure S4**. **System magnification characterization**. Profiles of 100 nm beads that were imaged at different stage position i) X = 0 µm, ii) X = 10 µm, iii) X = 20 µm. The resulting pixel size and magnification are 10 µm /(95-52)= 0.233 µm and 6.5 µm/0.233 µm = 27.9 x, which agrees with the theoretical magnification determined by the effective focal lengths of the objective and the tube lens (200 mm/7.2 mm = 27.8x)

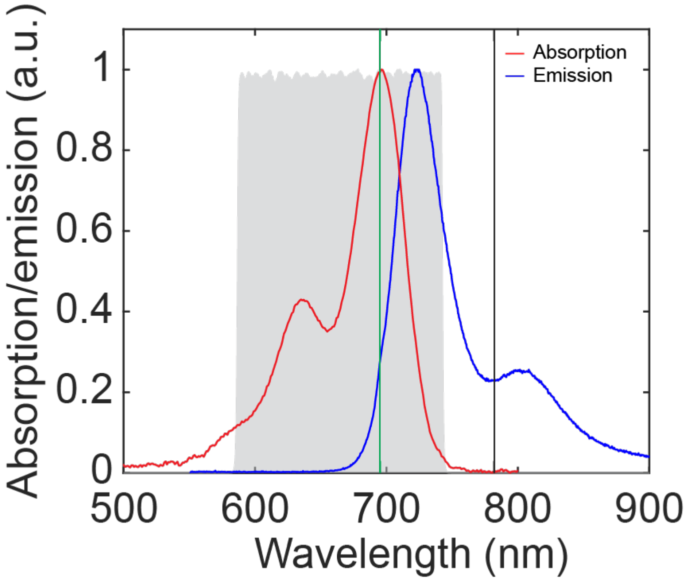


**Figure S5**. **Absorption and emission spectra of Rh800**. The gray indicates collection window, whereas the green and black lines indicate sum frequency (MIR+NIR) and NIR probe wavelength, respectively.

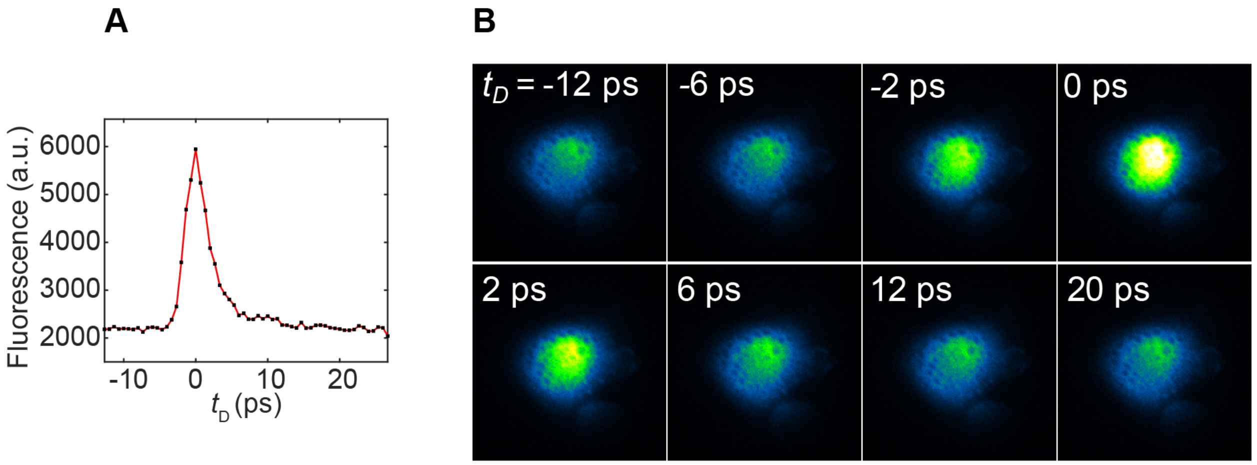


**Figure S6**. **Temporal profile characterization of Rh800 polymer sample** (A) Temporal profile characterization of Rh800 stained polymer sample (same as shown in Figure 2A). (B) Fluorescence images corresponding to different temporal delays $t_D$ between the MIR and the NIR probe lasers ($t_D$ >0 or $t_D$ < 0 indicates MIR or NIR probe pulses arriving first on sample, respectively; $t_D$ =0 indicates both laser pulses arriving the sample simultaneously).

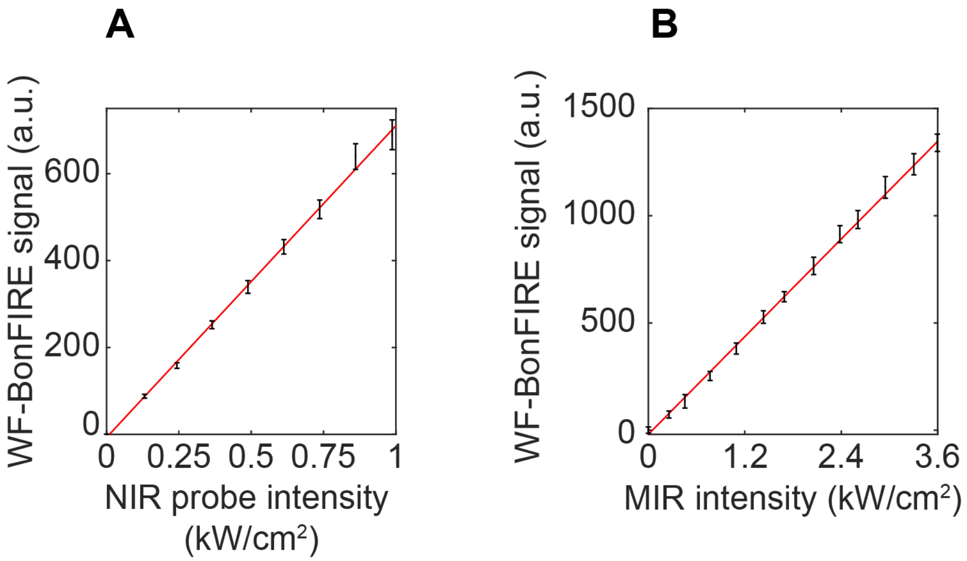


**Figure S7**. **Power dependence of WF-BonFIRE.** WF-BonFIRE signal dependence on the NIR probe power (A, MIR power was fixed at 4mW); and on the MIR power (B, NIR power was fixed at 466 mW).

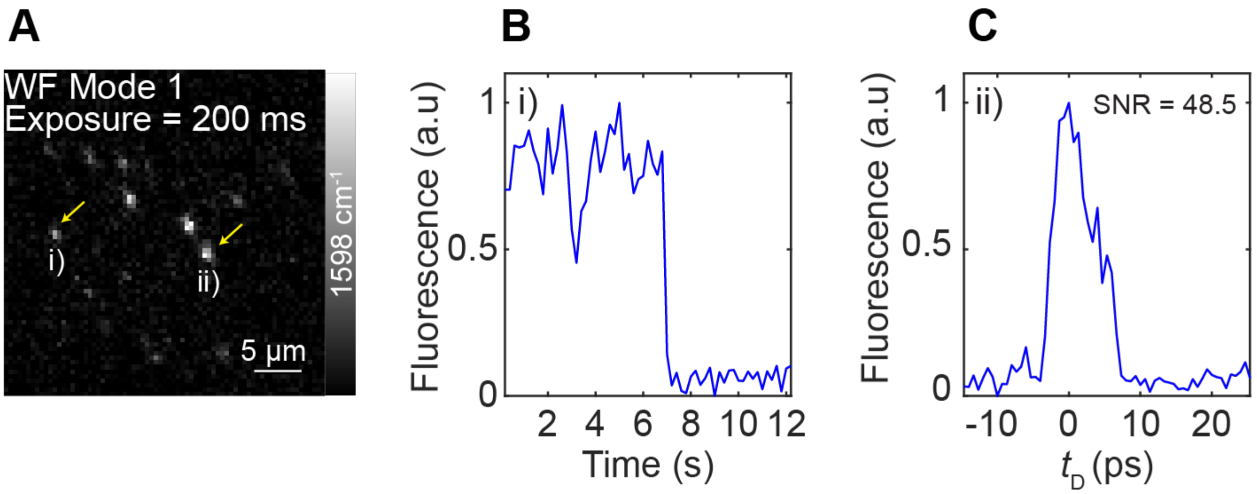


**Figure S8. Single-molecule WF-BonFIRE image at 200 ms exposure time.** (A) WF-BonFIRE image of single-molecule sample at 200 ms exposure time (same as shown in Figure 2D). (B) Single step photobleaching curve of molecule i) (yellow-arrowed in (A)). (C) Temporal sweep of molecule ii) (yellow-arrowed in (A)) for SNR characterization.

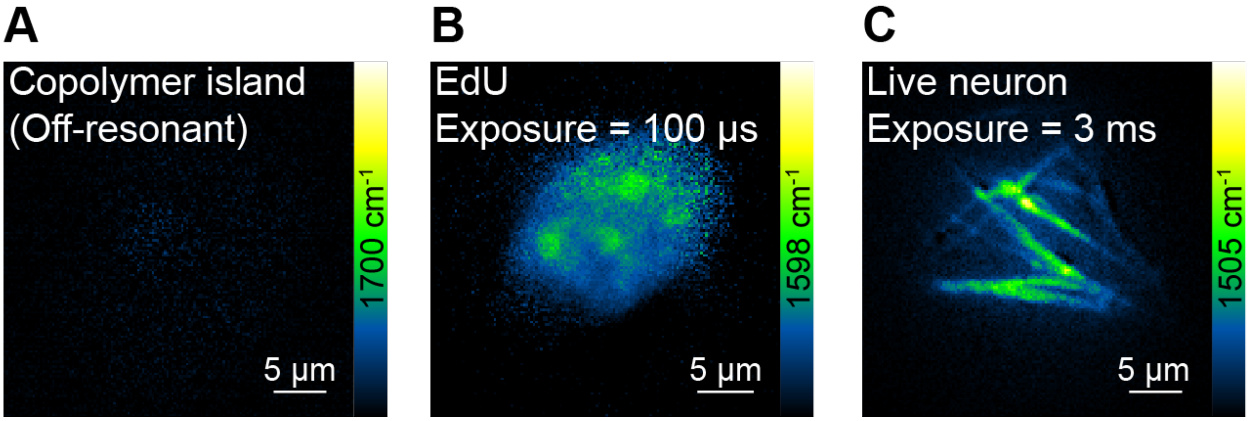


**Figure S9**. **WF-BonFIRE imaging (Mode 1) of various samples.** (A) WF-BonFIRE image of Rh800 copolymer island at off-resonance (1700 $cm^{-1}$). Exposure time: 17.6 µs. (B-C) WF-BonFIRE image targeting C=C vibration in ATTO680-click-labelled EdU in the nuclei of HeLa cells (B, 1598 $cm^{-1}$. Exposure time: 100 µs) and in Rh800-labelled mitochondria in live mouse neuronal cultures (C, 1505 $cm^{-1}$. Exposure time: 3 ms).

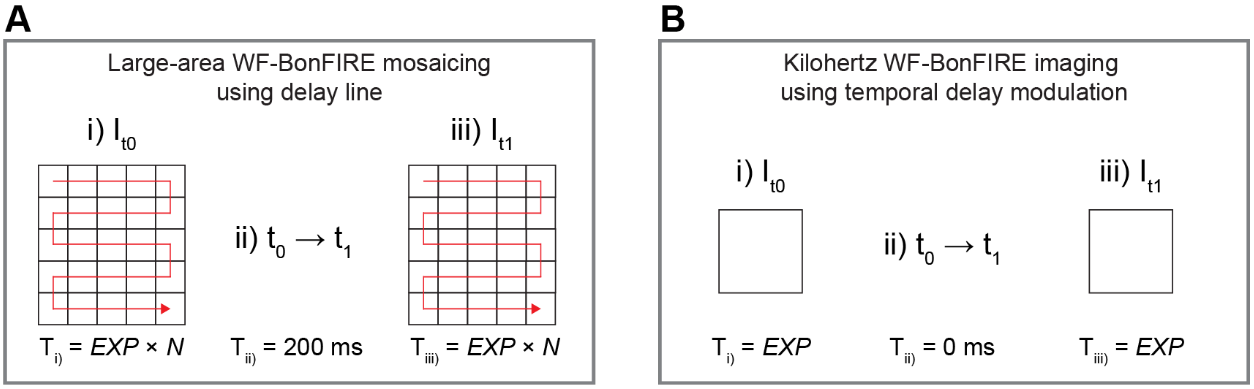


**Figure S10**. **Illustration for large-area mosaic WF-BonFIRE and kilohertz WF-BonFIRE using temporal delay modulation.** (A) Large-area WF-BonFIRE mosaicking using delay line. (B) Kilohertz WF-BonFIRE imaging using temporal delay modulation. *N* is the number of FOVs acquired in a mosaic. $T_{i)}$ and $T_{iii)}$ are the image acquisition time for the designated FOV when $t_D$ =0 and $t_D$ =20 ps. $T_{ii)}$ in (A) designates the delay line stabilization time, while in (B) is 0 ms due to the implementation of the temporal delay modulation scheme.

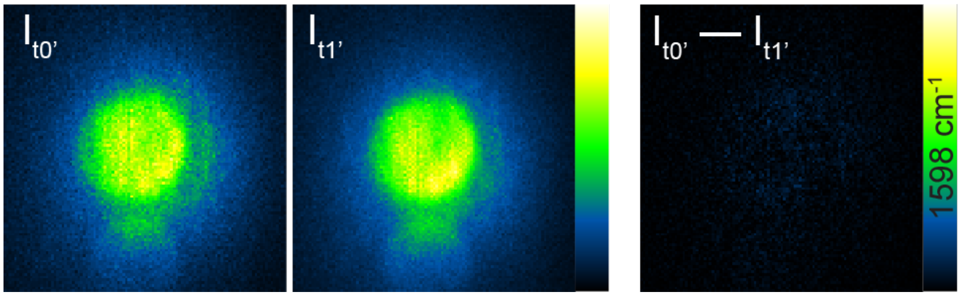


**Figure S11**. **Alignment characterization for the temporal delay modulation scheme**. WF-BonFIRE image ($I_{t0'} - I_{t1'}$) was generated by subtracting the subsequent camera frame from Path 1 ($I_{t0'}$) and Path 2 ($I_{t1'}$), where both temporal delays of $t_{0'}$ and $t_{1'}$ indicate when MIR and NIR pulses are temporally misaligned. The resulting image ($I_{t0'} - I_{t1'}$) from subtraction is a dark image with close to 0 intensities, indicating the absence of artifacts, ensuring the validity of the temporal modulation setup and alignment. FPS, 300 Hz, exposure, 1 ms.

**Movie S1. WF-BonFIRE imaging capturing Brownian motion in Rh800-stained E. coli at 150 frames per second (FPS) using temporal delay modulation.**
